# Development of a vanillate biosensor for the vanillin biosynthesis pathway in *E. coli*

**DOI:** 10.1101/375287

**Authors:** Aditya M. Kunjapur, Kristala L. J. Prather

## Abstract

Genetically encoded small molecule sensors can facilitate metabolic engineering by enabling high-throughput detection of metabolite concentrations, directed evolution of host and pathway enzymes, and dynamic regulation. The engineered *de novo* vanillin biosynthesis pathway assembled in *Escherichia coli* is industrially relevant and ideal for biosensor deployment given that the pathway requires only three heterologous enzyme-catalyzed reactions, generates naturally occurring metabolites, and may benefit from dynamic regulation. However, pathway flux is stalled and diverted by the activity of the *Homo sapiens* catechol O-methyltransferase, which is intended to catalyze the conversion of protocatechuate to vanillate. To confront this challenge, we constructed and applied a vanillate sensor based on the *Caulobacter crescentus* VanR-VanO system. Using components from a previously characterized *E. coli* promoter library, we achieved greater than 14-fold dynamic range in our best rationally constructed sensor. We characterized sensor substrate specificity and found that this construct and an evolved variant are remarkably selective, exhibiting no detectable response to the regioisomer byproduct isovanillate. We then harnessed the evolved biosensor to conduct rapid bioprospecting of natural catechol O-methyltransferases. We identified eight that appear to have greater desired activity than the originally used variant, including three previously uncharacterized O-methyltransferases. Collectively, these efforts enrich our knowledge of how biosensing can aid metabolic engineering and constitute the foundation for future improvements in vanillin pathway productivity.

## Introduction

Nature evolved a myriad of genetically encoded small molecule biosensors for detection and response to chemical stimuli. Allosteric transcription factors are a common form of biosensor and are commonly employed in inducible gene expression systems^1^ and in engineered genetic circuits^2^. The potential value of biosensors to metabolic engineers, who seek to maximize production of value-added products by manipulating cellular metabolism, is immense and multifaceted^3,4^. Genetically encoded biosensors facilitate metabolic engineering by enabling high-throughput detection of metabolite profiles, directed evolution of host and pathway enzymes^5–7^, and dynamic regulation^8–10^.

The *E. coli de novo* vanillin biosynthesis pathway^11,12^ is a highly industrially relevant pathway for which genetically encoded biosensors could be present for all three heterologous metabolites, and its potential as a testbed for multiplexed biosensing increases its academic relevance. Vanillin is the primary molecule responsible for vanilla flavor and is the largest flavor additive by volume, with an annual global market worth $650 million and a total volume of 18,000 tons^13^. It is also used as an intermediate for pharmaceuticals and specialty chemicals^14^. Notably, the vanillin biosynthesis pathway consists of three heterologous enzymes but numerous potential byproducts are also possible, which can be difficult to separate (Fig. 1). The enzyme AsbF from *Bacillus thuringiensis* catalyzes the conversion of an endogenous intermediate of the aromatic amino acid biosynthesis pathway, 3-dehydroshikimate, to protocatechuate (also referred to as 3,4-dihydroxybenzoate)^15^. The soluble catechol O-methyl-transferase from *Homo sapiens* (OMT_*Hs*_) catalyzes the next desired conversion of protocatechuate to vanillate^16–18^. Finally, the carboxylic acid reductase from *Nocardia iowensis* (Car_*Ni*_) catalyzes the final desired conversion of vanillate to vanillin^19,20^. Use of an engineered *E. coli* host (the RARE strain) minimizes rapid endogenous conversion of vanillin to vanillyl alcohol^11^. However, other products are commonly formed downstream of the protocatechuate branch point due to the broad substrate specificity of OMT_*Hs*_ and Car_*Ni*_. OMT_*Hs*_ catalyzes the formation of isovanillate by transferring a methyl group to the hydroxy group at the para rather than meta position. Car_*Ni*_,· can also convert protocatechuate to protocatechualdehyde, which is a toxic intermediate, though it is possible for OMT_*Hs*_ to subsequently convert protocatechualdehyde into vanillin.

**Figure 1.**
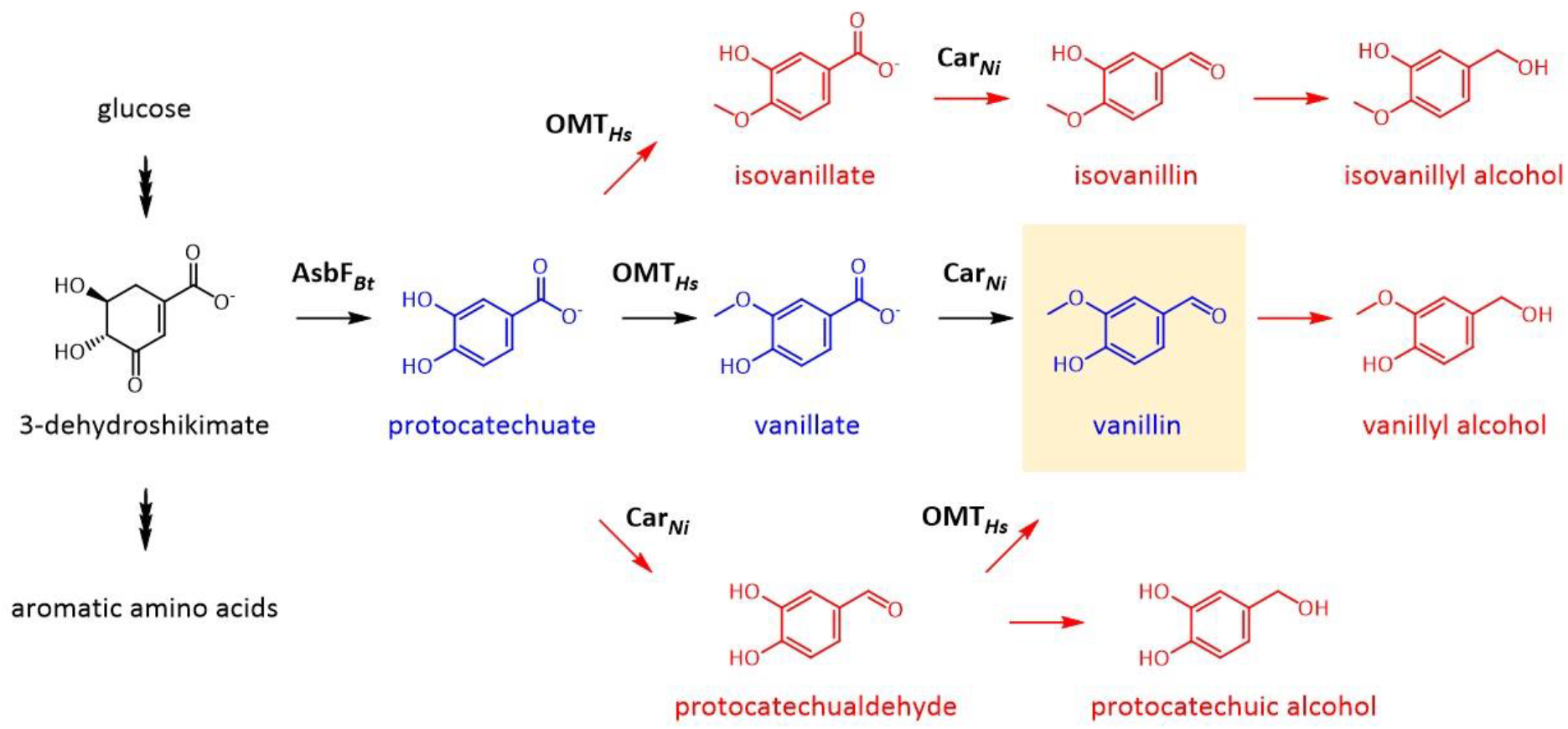
Metabolic pathway diagram of the engineered *de novo* vanillin biosynthesis pathway and potential byproducts in *E. coli.* Heterologous steps begin with AsbF-catalyzed conversion of endogenous 3-dehydroshikimate into protocatechuate. Protocatechuate, vanillate, and vanillin form desired heterologous metabolites and are shown in blue. Undesired metabolites that form due to low specificity of Car and OMT enzymes or due to endogenous aldehyde reductase activity in wild-type *E. coli* strains are shown in red.

The ability to sense vanillate could improve vanillin titers through several means. Based on the observation that protocatechuate accumulates, we previously found that OMT_*Hs*_ steadily loses activity during the first 48 hours after induction and that the co-factor *S*-adenosyl-methionine (SAM), which is the required methyl donor, appears to become scarce as fermentation time proceeds^12^. A vanillate sensor could be used to improve OMT_*Hs*_ activity, to engineer a host for increased SAM pool size, or to rapidly screen the natural diversity of OMTs in search of a better variant to relieve this pathway bottleneck. In principle, a vanillate sensor could also be used to delay expression of CarN· until vanillate accumulated, which should favor flux through desired heterologous reaction steps and decrease formation of toxic protocatechualdehyde.

We examined the literature for allosteric transcription factors that respond to the three desired heterologous metabolites - protocatechuate, vanillate, and vanillin - given that they are naturally obtained through lignin degradation (Fig. 1B). An *E. coli* sensor for protocatechuate was previously constructed by importing the PcaU activator and associated promoter sequence from *Acinetobacter^21^.* This sensor was reported to exhibit a graded (or analog) response across millimolar concentrations of protocatechuate, with high substrate specificity and approximately 10-fold dynamic range. In addition to the protocatechuate sensor, multiple efforts have been undertaken to develop an *E. coli* sensor specific for vanillin. The QacR transcriptional repressor was computationally redesigned to bind to the non-native substrate vanillin and led to variants that exhibited dose-dependent de-repression *in vitro* and *in vivo^22^.* Additionally, recent work has revealed that a native *E. coli* promoter may be responsive specifically to vanillin^23^. However, we found no dedicated effort to construct and characterize a sensor responsive to vanillate in *E. coli.*

Here, we report the development, characterization, and application of a genetically encoded sensor for vanillate in *E. coli* based on the *Caulobacter crescentus* VanR-VanO system. Our initial results obtained through rational modifications demonstrate the respective influence of repressor promoter strength and operator site positioning on sensor dynamic range. Our subsequent results showcase the exquisite molecular specificity of the VanR transcription factor through the dose-responses obtained for original and evolved vanillate biosensor constructs. We conclude by illustrating the value of biosensor-based bioprospecting by screening the natural diversity of catechol *O*-methyltransferases sampled across all domains of life. Our approach rapidly identifies variants that appear to have higher desired activity than OMT_*Hs*_, including three previously uncharacterized *O*-methyltransferases.

## Results and Discussion

Vanillate-responsive transcriptional repressors (VanR) exist in several bacteria, including *Äcinetobacter^24^, Caulobacter^25^, Corynebacterium^26^, Myxococcus^27^,* and *Pseudomonas^28^.* These VanR variants bind to different operator (VanO) sites consisting of inverted repeats ranging from AACTAACTAA(N4)TTAGGTATTT in *Cornyebacterium glutamicum* to ATTGGATCCAAT in *Caulobacter crescentus* (Fig. 2A/B). We sought to import the *C. crescentus* system into *E. coli* given the relatively small operator site and because a vanillate-inducible expression system was successfully developed for use in *C. crescentus^25^.* At first, we considered the three base pair overlap of the natural VanO sequence with the *C. crescentus* −10 site a potential obstacle to direct promoter transfer and sought to test rational alternatives for *E. coli* (Fig. 2C). Placement of the VanO site at the same position in relation to the *E. coli* - 10 site (leading to TATATT instead of the consensus TATAAT^29^) is predicted to significantly disfavor interaction with the *E. coli* sigma factor 70^30^.

**Figure 2.**
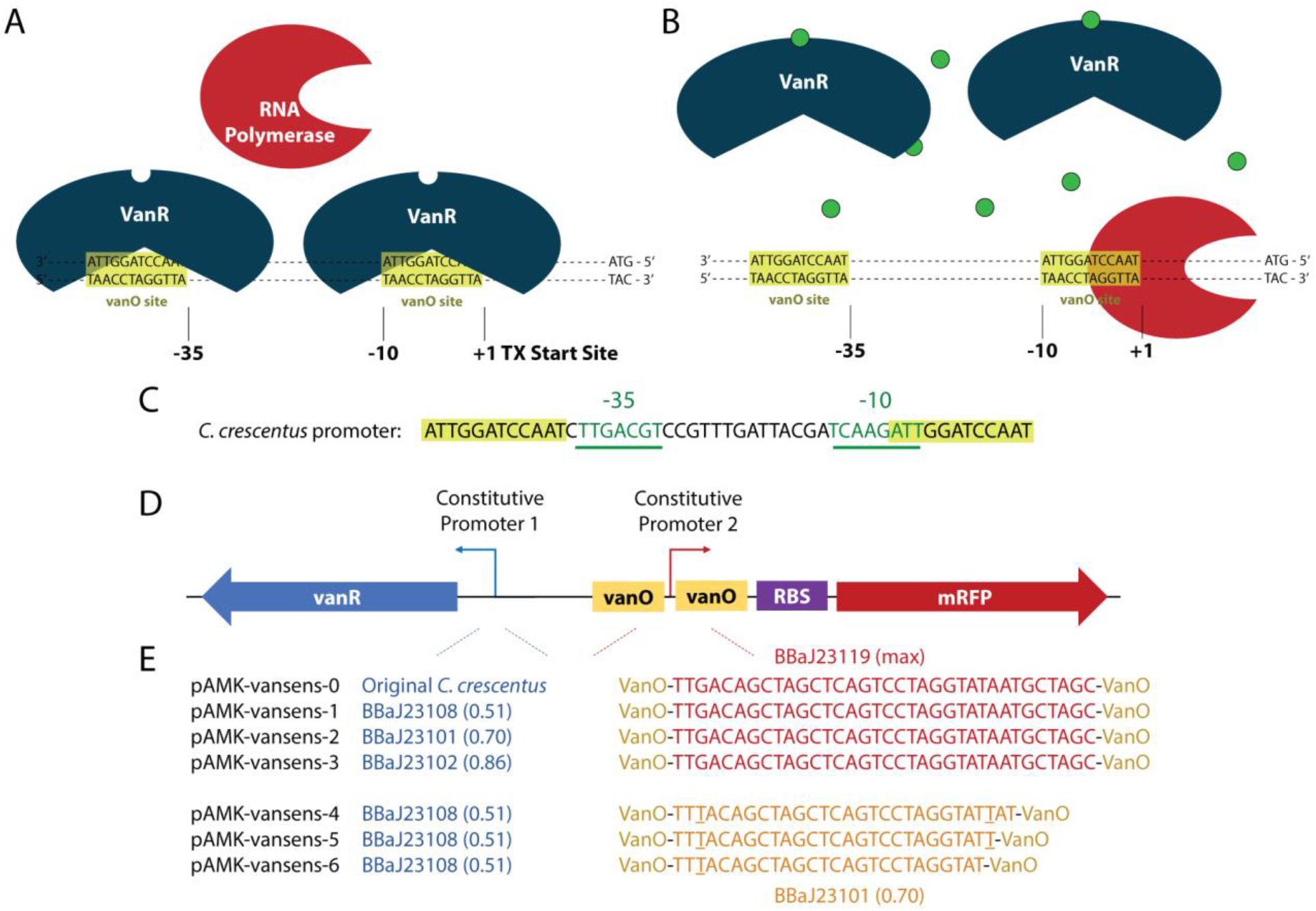
Original vanillate biosensor components and design. (A) Cartoon illustrating desired binding of vanillate repressor VanR to vanillate operator VanO sites. VanO sites are adjacent to −35 and −10 regions upstream of transcriptional start site, thereby hindering binding of sigma factor and RNA polymerase complex. (B) Cartoon illustrating desired unbinding of allosterically regulated VanR from VanO in the presence of vanillate, thereby enabling transcription. (C) Sequence of the native pVanAB promoter regulated by VanR in *Caulobacter crescentus,* showing that one of the two VanO sites overlaps with the −10 region. The consensus −10 region in *E. coli* is TATAAT and the consensus −35 is TTGACA. (D) High-level abstraction of vanillate sensor design. (E) Candidate promoter sequences tested experimentally in this manuscript. The two underlined “T”s in BBaJ23101 reflect the sequence difference between it and BBaJ23119.

We aimed to construct a functional vanillate sensor in *E. coli* using the *C. crescentus* VanR-VanO system to regulate expression of a red fluorescent protein (mRFP) (Fig. 2D). The complete list of candidate promoter sequences used for expression of codon-optimized VanR and mRFP are shown in Fig. 2E. Our initial vanillate sensor construct (pAMK-vansens-0) used the native *C. crescentus* promoter upstream of VanR to drive its expression. A strong Anderson promoter flanked by VanO sequences was placed upstream of mRFP. *E. coli* harboring this construct showed little to no change in mRFP fluorescence upon addition of 1 mM vanillate. As we began to increase expression of VanR using Anderson library promoters instead of the native *C. crescentus* promoter (previously valued at 0.51/0.70/0.86 expression strength – pAMK-vansens-1/2/3, respectively), we observed an increased ratio of fluorescence in the presence of vanillate to fluorescence in the absence in the absence of vanillate (Fig. 3A). However, we observed substantial leaky fluorescence (Fig. 3A). After realizing that the downstream VanO site was neither adjacent to nor overlapping with the −10 site, we generated three additional constructs with the VanO site placed adjacent (pAMK-vansens-4) or with 1 or 2 bp overlap (pAMK-vansens-5 or pAMK-vansens-6, respectively). *E. coli* harboring these constructs exhibited low fluorescence in the absence of vanillate and increasing dynamic range as overlap of VanO and −10 site increased (Fig. 3B). The best performing construct, pAMK-vansens-6, exhibited a dynamic range of approximately 14-fold and was chosen for further characterization.

**Figure 3.**
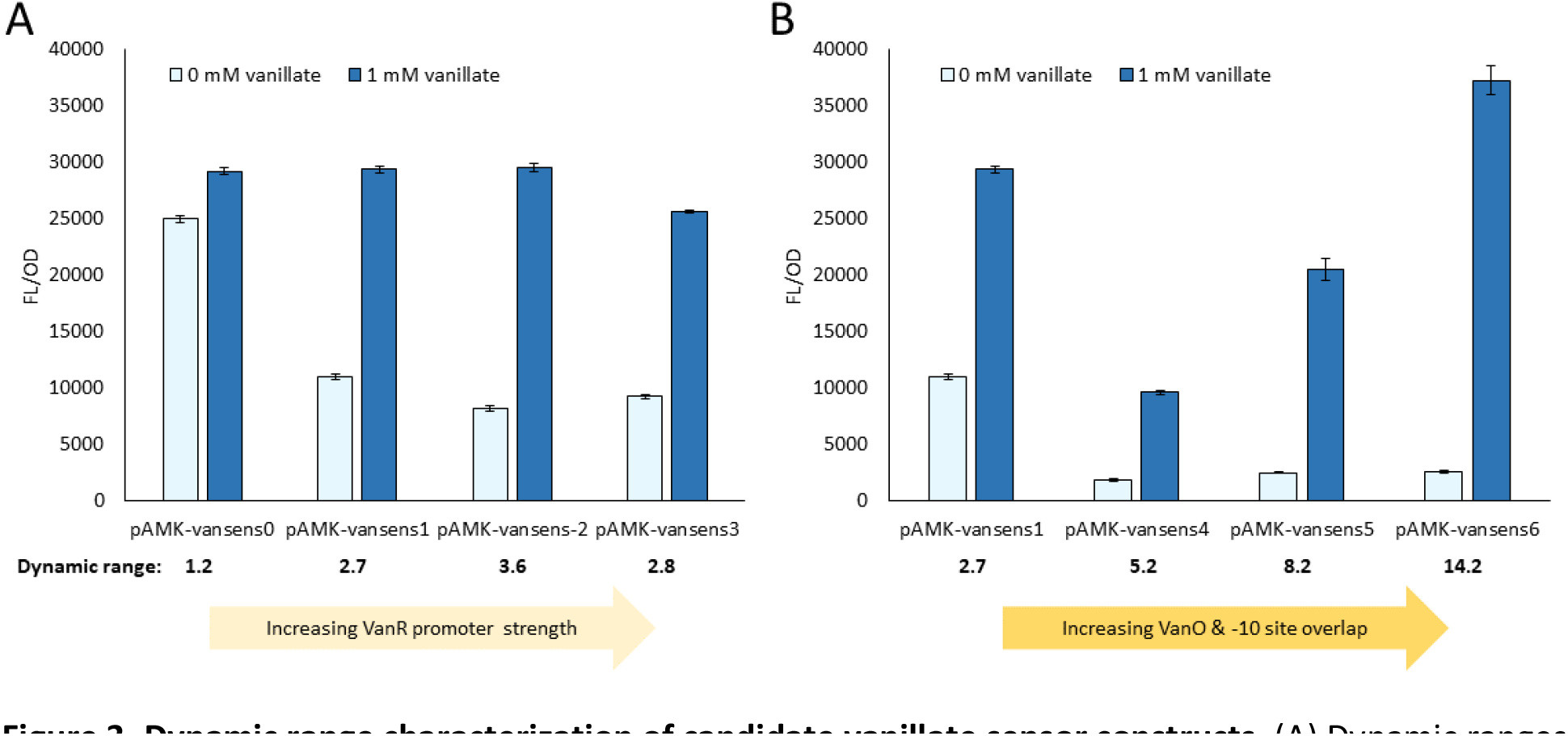
Dynamic range characterization of candidate vanillate sensor constructs. (A) Dynamic ranges observed using candidate vanillate sensor constructs expressing different VanR promoter strengths. (B) Dynamic ranges observed using constructs containing different extents of overlap between VanO and *E. coli* −10 site.

We characterized the dose-response behavior of this vanillate sensor for five relevant substrates that are typically produced after co-expression of vanillin pathway genes in *E. coli:* protocatechuate, vanillate, isovanillate, protocatechualdehyde, and vanillin. These substrates were provided at concentrations ranging from 0.01 mM to 1 mM, the latter of which is in the order of magnitude of observed accumulation of protocatechuate and vanillate in the absence of Car_*Ni*_ expression^31^. Furthermore, we compared vanillate sensor performance with the best available sensor reported for protocatechuate as a reference and to explore future potential for dual sensing. We cloned the published protocatechuate sensor (PcaU and promoter region) upstream of mRFP in place of equivalent components in the vanillate sensor to create an otherwise identical construct (pAMK-protosens-1). Based on an endpoint assay 24 hours after substrate supplementation, we observed graded dose-response to protocatechuate consistent with the literature^21^ (Fig. 4A). Unfortunately, the protocatechuate sensor also activated in the presence of vanillate, which had been reported. In contrast, our best vanillate sensor (pAMK-vansens-6) exhibited high sensitivity to vanillate, with half-maximal activation occurring below 10 μM and full activation below 100 μM range (Fig. 4B). The high sensitivity and more quickly saturating response were concerning given our aspirations to ascertain relative concentration resulting from biosynthesis in different strains rather than merely whether vanillate was present. Our concerns were mitigated by the excellent specificity of the vanillate sensor, which exhibited minimal response to protocatechuate and remarkably no response to the regioisomer isovanillate. Many catechol *O*-methyltransferases, including OMT_*Hs*_, possess the undesired capacity to methylate a hydroxy group on the para rather than meta position of the aromatic ring relative to the carboxylic acid group, which gives rise to isovanillate from protocatechuate^17,32^. Isovanillate is particularly problematic given that *CarNi* will convert it to isovanillin, and these two regioisomers add hurdles to separation from vanillate and vanillin for accurate quantification or product purification.

**Figure 4.**
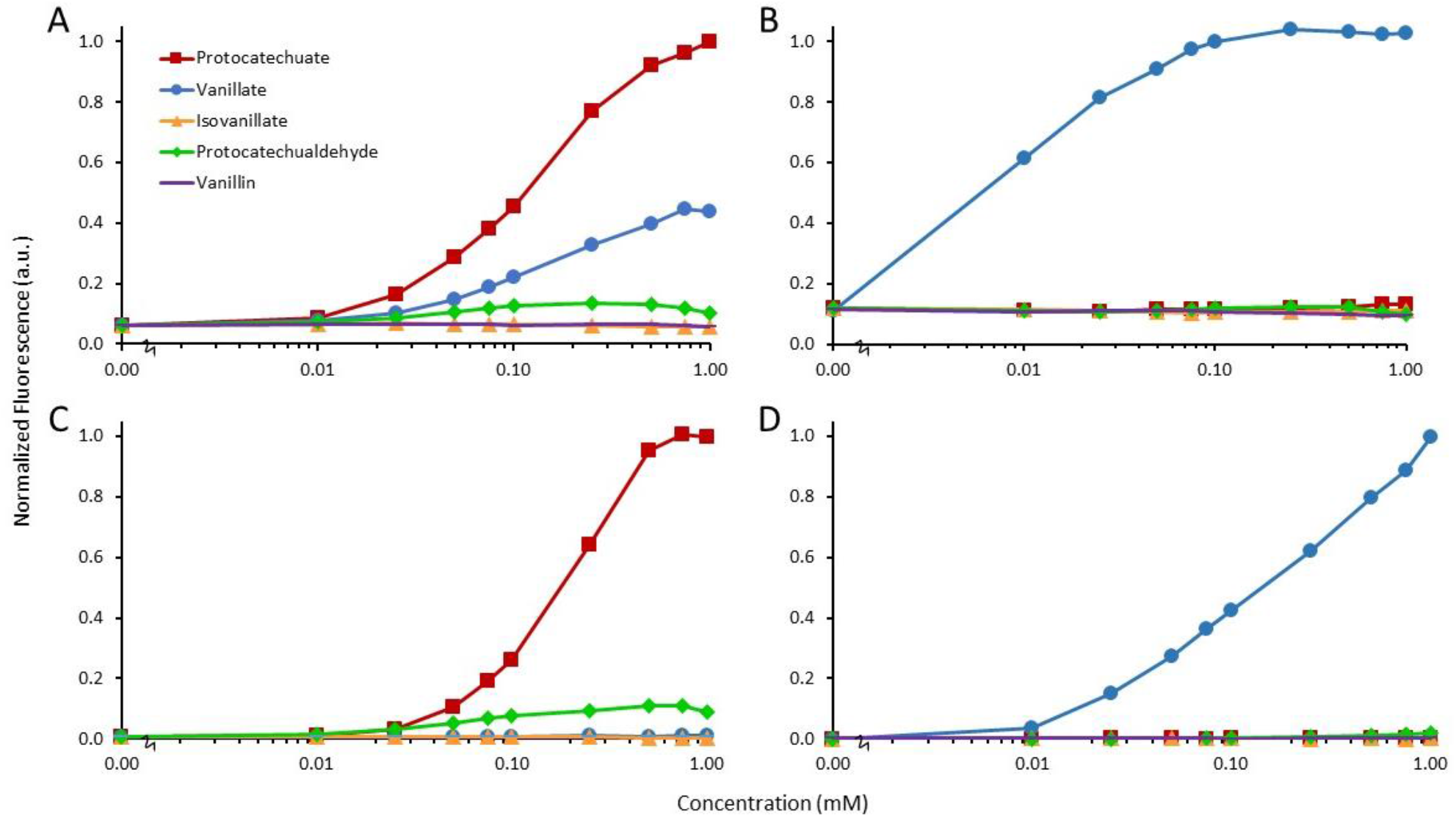
Dose-response characterization of original and evolved sensors intended for protocatechuate and vanillate, respectively. (A) Original protocatechuate sensor (from Ref. 21 and cloned into pAMK030). (B) Original vanillate sensor (pAMK022). (C) Evolved protocatechuate sensor (pAJM690, from Ref. 34). (D) Evolved vanillate sensor (pAJM773, from Ref. 34).

We next tested the vanillate sensor in cells that expressed the vanillin biosynthesis pathway. *E. coli* RARE cells harboring pathway constructs and pAMK-vansens-6 exhibited a low level of fluorescence in LB medium and several fold additional activation in response to externally supplied vanillate (SI Fig. 1A). These cells also demonstrated endogenous activation of the vanillate sensor in M9 minimal media supplemented with glucose and IPTG for induction of pathway genes (SI Fig. 1B). However, the sensor exhibited full activation under this condition and full activation when no IPTG was added, suggesting that the sensor is saturated by leaky uninduced pathway expression in the presence of glucose.

To overcome these challenges, we pursued further engineering of our vanillate sensor. We provided colleagues in the laboratory of Professor Christopher Voigt (MIT Department of Biological Engineering, Cambridge, MA) this sensor as well as the recloned protocatechuate sensor for evolution by compartmentalized partnered replication^33^, which was successfully used to improve the dynamic range of these two sensors as well as ten others as very recently reported^34^. As before, we investigated the dose-response and substrate specificity of these evolved sensors, which drive expression of YFP instead of mRFP (Fig. 4C and 4D). We observed improvements in dynamic range and protocatechuate sensor specificity consistent with the literature^34^. Fortunately, the evolved vanillate sensor, which includes mutations to VanR, maintains an absence of response to isovanillate. Although the evolved vanillate sensor was originally found to have increased cooperativity relative to the original vanillate sensor, we found over repeatable experiments in our hands that the dose-response of the evolved vanillate sensor was more graded than the original vanillate sensor and that it saturated at a higher vanillate concentration. Both features increase biosensing efficacy for metabolic engineering contexts. Th discrepancy may be a function of media conditions, supplementation time, and/or assay endpoint time.

Given that the evolved vanillate sensor can generate proportional fluorescence at sub-millimolar vanillate concentrations, has extremely low background fluorescence, and does not respond to related vanillin pathway molecules including isovanillate, we harnessed it to improve the vanillin pathway by biosensor-based bioprospecting. As discussed earlier, we previously identified OMT_*Hs*_ as the bottleneck for pathway flux and hypothesized that other *O*-methyltransferases may be better at catalyzing vanillate formation, especially in a bacterial host. As a starting point for sampling the natural diversity of *O*-methyltransferases, we considered all relevant literature including published patent applications. One patent application filed by Evolva S.A. and International Flavors and Fragrances, Inc. (IFF) describes the testing of different *O*-methyltransferases expressed in yeast and displays data for roughly 20 natural variants^32^. We compiled a shortlist consisting of the best *O*-methyltransferases reported and then used protein BLAST on the NCBI nucleotide collection to identify putative enzymes and other *O*-methyltransferases that had not been previously tested. We sampled across the tree of life but biased our final candidate list of sixteen towards bacterial, fungal, and archaeal sources given our bacterial host (Fig. 5).

**Figure 5.**
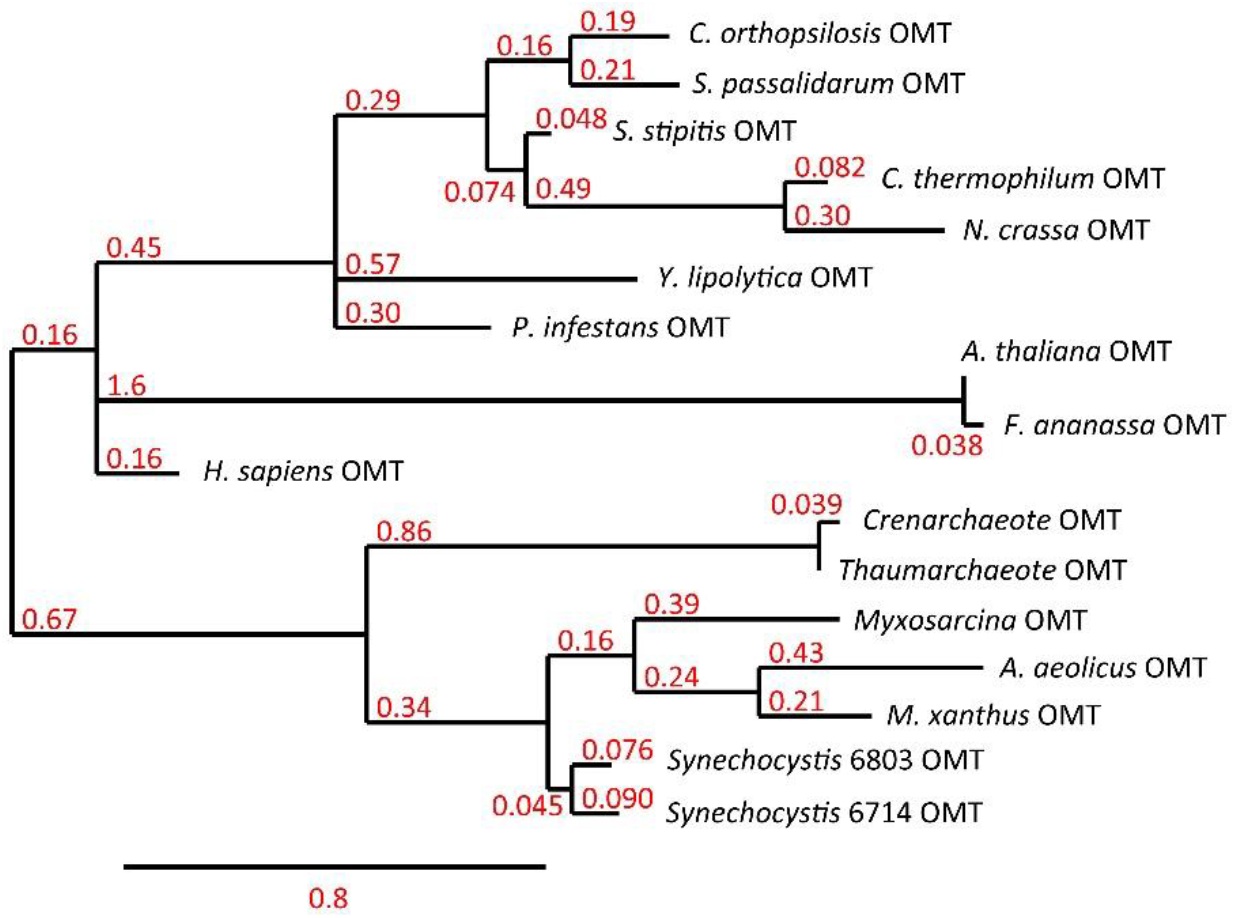
Phylogenetic tree of naturally occurring candidate OMT variants for biosensor-based bioprospecting. Sixteen variants from diverse sources across the tree of life were chosen for rapid screening of *in vivo* O-methyltransferase activity on exogenously supplemented protocatechuate. All are distant from the originally characterized *H. sapiens* OMT. Phylogenetic tree was generated using Phylogeny.fr and protein sequence inputs. Red values indicate branch lengths.

We screened these *O*-methyltransferases by co-transforming constructs harboring the evolved vanillate sensor and constructs harboring each of the library members into the *E. coli* RARE *ΔmetJ* strain^12^. The deletion of the MetJ regulator of methionine biosynthesis is one of several modifications previously shown to improve conversion of protocatechuate to vanillate, presumably due to increased SAM pool size. We tested these cultures with IPTG supplementation at inoculation, which induces O-methyltransferase expression, and with or without 4 mM protocatechuate supplementation. We then measured final OD and normalized fluorescence (YFP/OD) 24 hours after inoculation. Our first observation was that one of the variants grew substantially more slowly than the rest and achieved a lower final OD by approximately two-to three-fold (Fig. 6A). This variant, the O-methyltransferase from *Phytophthora infestans* or OMT_*Pi*_, also exhibited the highest normalized fluorescence despite its potential toxicity. Because the expression of OMT_*Pi*_ resulted in a lower final OD independent of exogenous protocatechuate provision, the toxicity of OMT_*Pi*_ expression may be unrelated to its high enzymatic activity or may be due to activity on other endogenous substrates. Given its adverse effect *in vivo,* this enzyme may be better suited to *in vitro* strategies for vanillin biosynthesis^35^ if high protein yields can be obtained.

**Figure 6.**
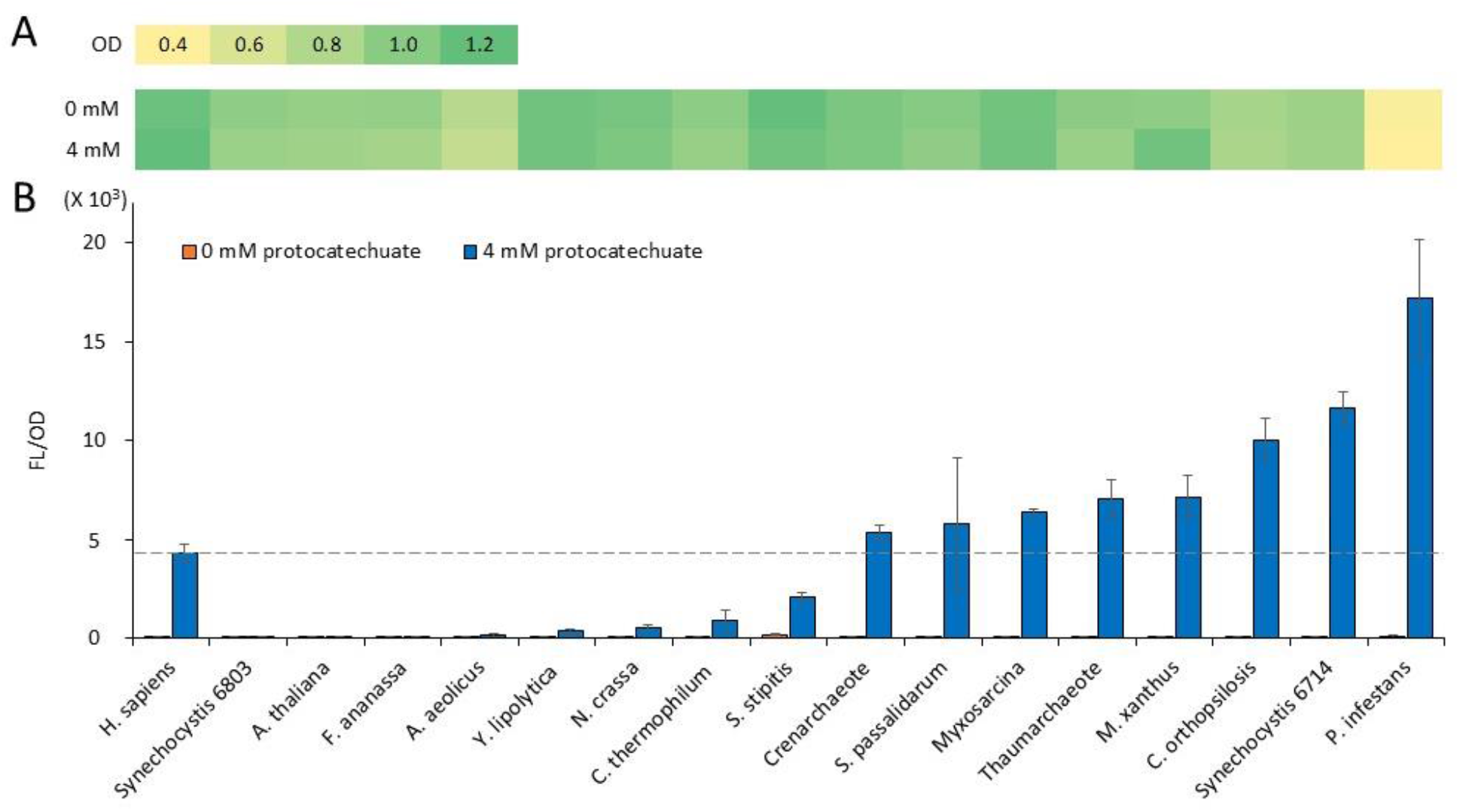
Biosensor-based bioprospecting of naturally occurring O-methyltransferase variants. (A)Average final optical densities (OD) of cultures that harbor evolved vanillate biosensor and that express OMT variants in the absence (0 mM) or presence (4 mM) of exogenously supplied protocatechuate at the 24h endpoint of the assay. Cultures that express *P. infestans* OMT achieved lower final ODs. (B) Endpoint OD-normalized fluorescence of cultures that harbor vanillate biosensor and that express OMT variants in the absence or presence of exogenously supplied protocatechuate. Dashed line demarcates performance of existing OMT to facilitate identification of improved variants.

The results of our screen suggest that eight *O*-methyltransferases could have greater desired *in vivo* activity on protocatechuate when expressed in *E. coli* compared to our starting OMT_*Hs*_, and that three previously uncharacterized *O*-methyltransferases may have utility for this pathway and for broader biotechnology applications. Of the six best performing OMTs screened, the variants from *Phytophthora infestans* (top performing) and *Candida orthopsilosis* (third best performing) are described in the patent application filed by Evolva and IFF^32^. The second best performing OMT is from *Synechocystis* sp. PCC 6714 and is previously uncharacterized to the best of our knowledge. A report describing the draft genome sequence of PCC 6714 identified a genomic island containing a putative methyltransferase upon comparative genome analysis of the related *Synechocystis* sp. PCC 6803 strain^36^. The complete genome sequence record for PCC 6714 contains gene annotation of “Predicted O-methyltransferase YrrM” based on function prediction from protein sequence similarity^37^. Our finding that the PCC 6714 variant is so active is particularly interesting given the lack of activity observed from the PCC 6803 variant, which shares 86% protein sequence identity. When previously expressed in E. coli, the PCC 6803 variant exhibited preference for trihydroxylated flavones and reduced affinity for protocatechuate (kcat/KM = 37 M^−1^ s^−1^)^38^. *Myxococcus xanthus* is the source of our fourth best performing OMT, which is known to prefer protocatechuate as a substrate (kcat/KM = 5.5 × 10^3^ M^−1^ s^−1^)^39^ and to have high regioselectivity for the meta position of protocatechuate despite having high regioselectivity for the para position of other substrates^40,41^. Like the OMT from PCC 6714, the fifth and sixth best performing OMTs, which are from uncultured marine thaumarchaeote KM3_66_E12^42^ and from Myxosarcina sp. GI1 respectively, have not been characterized and are annotated as “Predicted O-methyltransferase YrrM”.

Overall, this work seeks to overcome a critical limitation of the industrially relevant vanillin biosynthesis pathway through the development, characterization, and application of a genetically encoded biosensor for the heterologous intermediate vanillate. Biosensing in *E. coli* was successfully achieved by importing transcriptional regulator and operator components from *C. crescentus* and by introducing rational modifications in promoter sequences. The vanillate biosensor was then further evolved by directed evolution, and these biosensors along with similar biosensors for protocatechuate were evaluated for their substrate specificity and dose-response. Three of these sensors – the original vanillate sensor and the evolved vanillate and protocatechuate sensors – each exhibited sufficiently high specificity to enable their future use for improving the vanillin pathway. We demonstrated the value of the vanillate biosensor by using it to rapidly bioprospect alternative *O*-methyltransferases and our screen suggests that several variants will outperform our original choice, including three previously uncharacterized proteins. The effort described here lays the foundation for subsequent vanillin pathway improvements as well as the ability to investigate the value of multiplexed biosensing of all desired heterologous metabolites in the vanillin pathway.

## Methods

### Strains and Plasmids

*E. coli* strains and plasmids used in this study are listed in SI Table 1. Molecular biology techniques were performed according to standard practices^43^ unless otherwise stated. Molecular cloning, vector propagation, and the majority of biosensor characterization were performed in DH5α. The host strain used for vanillin biosynthetic pathway co-expression experiments was the *E. coli* RARE strain (Addgene Catalog #61440)^11^, which is derived from E. coli K-12 MG1655(DE3). Oligonucleotides were purchased from Sigma and are listed in SI Table 2. Q5 High Fidelity DNA Polymerase (New England Biolabs, MA) was used for DNA amplification.

The VanR and PcaU genes were synthesized as codon-optimized gBlocks (Integrated DNA Technologies, CA) and their sequences are included in SI Table 3. Original sensor constructs were cloned into the Duet vector system (Novagen, WI) using restriction digest-based cloning. Restriction enzymes and T4 DNA ligase were purchased from New England Biolabs. Propagated constructs were purified using a QIAprep Miniprep Kit (Qiagen, CA) and agarose gel fragments were purified using a Zymoclean Gel DNA Recovery Kit (Zymo Research, CA). All constructs were confirmed to be correct by nucleotide sequencing (Genewiz, NJ).

Constructs for biosensor-based screening of natural O-methyltransferase diversity were generated using Gibson assembly^44^ from homemade assembly mixtures (enzymes from New England Biolabs, NTPs and other chemicals from Sigma). O-methyltransferase protein sequence accession numbers and corresponding codon-optimized nucleotide sequences that were ordered as IDT gBlocks are also included in SI Table 3.

Additional description of methods (Chemicals, Culture conditions, Fluorescence measurements, and Metabolite analysis) can be found in Supporting Information.

## Acknowledgements

We acknowledge Professor George Church (Harvard Medical School, Boston, MA) for limited use of a fluorescent 96-well plate reader (Biotek H4 Synergy) and Spencer Wenck for assistance in constructing the original vanillate sensor. We are also grateful to Professor Chris Voigt (MIT, Cambridge, MA) for sensor evolution and for sharing evolved sensor constructs. This research was initially supported by the National Science Foundation through the Synthetic Biology Engineering Research Center (Synberc, Grant No. EEC-0540879) and through a Graduate Research Fellowship to AMK. O-methyltransferase sequences were supported by a grant from the MIT-Portugal Program (Grant Number 6937814).

## Supporting Information

Supporting Information is available free of charge.

Supporting Information (PDF) contains additional description of methods, a figure demonstrating sensor activation during pathway co-expression, and tables of plasmids, oligonucleotides, and synthetic gene (gBlock) sequences.

